# Coaxial line-scanning Brillouin microscopy

**DOI:** 10.1101/2025.02.25.640150

**Authors:** Chenjun Shi, Jitao Zhang

**Affiliations:** Biomedical Engineering Department, Institute for Quantitative Health Science & Engineering, Michigan State University, MI 48824, USA

## Abstract

Confocal Brillouin microscopy (CBM) enables high-resolution mechanical imaging but has slow acquisition speeds due to its point-by-point scanning strategy. Line-scanning Brillouin microscopy (LSBM) offers imaging acquisition speed improvements but faces challenges such as beam distortion in biaxial configurations and insufficient extinction ratio due to the single-stage VIPA spectrometer. To overcome these limitations, we developed a coaxial line-scanning Brillouin microscopy (cLSBM) system by using a two-stage parallel VIPA spectrometer. The coaxial design minimizes image distortion, and the two-stage parallel VIPA spectrometer significantly enhances the rejection of non-Brillouin noises. Experiment results showed that the first VIPA, served as a filter for noise rejection, has a rejection ability of 18 dB. The system was characterized by standard materials including ethanol and water, achieving a precision of 7.5 MHz and 12.6 MHz respectively. In the next step, we will optimize the system to further enhance noise rejection and utilize this setup to investigate the evolution of tissue mechanics during embryonic development.

## Introduction

Confocal Brillouin microscopy (CBM) is based on spontaneous Brillouin scattering, where the frequency shift of the scattered light is introduced due to photon-phonon interaction. CBM provides a mechanical imaging with diffraction-limited spatial resolution for various biological samples[1]. However, due to the low scattering efficiency of the spontaneous Brillouin scattering process and the point-by-point scanning strategy, the acquisition speed of CBM remains slow. To improve the speed, line-scanning Brillouin microscopy (LSBM), which illuminates samples with a needle-shape beam, provides a multiplexing solution to acquire Brillouin spectra of multiple points in a single shot[2, 3]. We have demonstrated that the LSBM can be 50-100 times faster than CBM for 2D/3D mechanical imaging of cellular spheroids. However, currently LSBM mainly uses biaxial configuration, in which the illumination axis and the detection axis are separated. Therefore, it suffers from beam distortion caused by the mismatch of the refractive index between the sample and the surrounding medium. In addition, the setup is built in the horizontal plane of an optical table, which requires the sample to be mounted in hydrogel and thus is not suitable for long-term live imaging. In addition, for most biological samples, the non-Brillouin noise is usually much stronger than the Brillouin signal, which requires the spectrometer to have a high extinction ratio. In CBM, it can be achieved by using a two-stage cross-axis VIPA design (Figure 1(a)), where the light can be dispersed twice in two orthogonal axes so that more noise can be rejected [4]. However, in the case of LSBM, the illumination line already occupies one spatial axis. Therefore, current line-scanning setup can only use one-stage VIPA configuration, which cannot provide enough extinction ratio to reject non-Brillouin noise. The current solution is to use the laser source with a specific wavelength and accompanied by a gas chamber to absorb the laser frequency (e.g., 780 nm laser with Rb gas chamber) [3]. Although effective, the current solution is limited to 780-nm laser and does not support other wavelengths.

**Figure 1.**
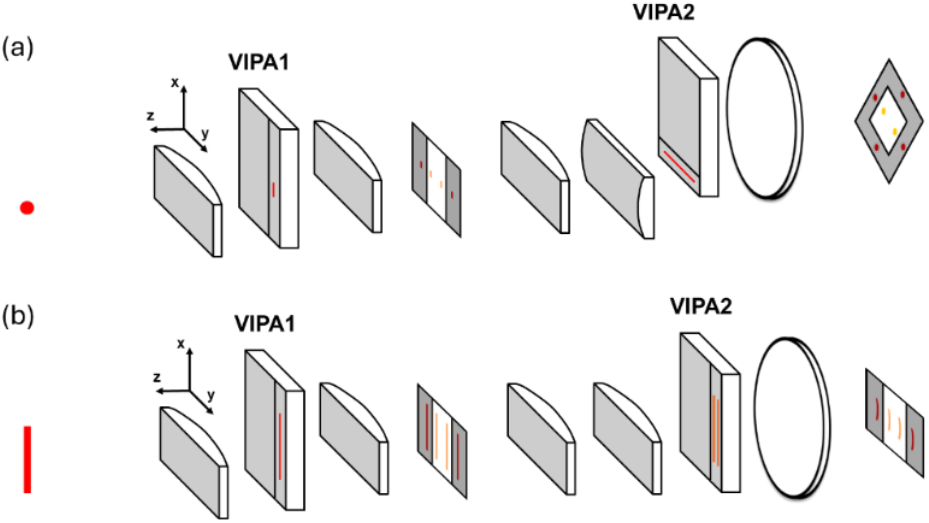
(a) two-stage cross-axis VIPA design for CBM, in which VIPA1 has a dispersion axis in the y direction and VIP2 has a dispersion axis in the x direction, and thus the final Brillouin pattern shows a dispersion in both x and y axis; (b) two-stage parallel VIPA design for line-scanning, in which both VIPA1 and VIPA2 have a dispersion axis on y direction, and thus the final Brillouin pattern shows a dispersion in y axis, while the other axis (x axis) is used as the spatial axis for the line-scanning setup.

To address these limitations, we developed a coaxial line-scanning Brillouin microscopy (cLSBM) by using a two-stage parallel VIPA spectrometer. In coaxial configuration, the illumination and detection axes share the same optical path. As a result, the cLSBM can reduce beam distortion and is thus suitable for optically heterogeneous samples. In addition, we adapted the setup in an inverted configuration so that samples can be imaged in their native culturing dishes without mounting, enabling longitudinal imaging of important biological events. Furthermore, in the spectrometer part, inspired by a two-stage parallel VIPA configuration in CBM, where the first VIPA was used as a filter for enhanced non-Brillouin noise rejection and the second VIPA was used as the spectrometer[5], we adapt this design to be available in line-scanning setup (Figure 1(b)). Take together, the system provided a higher extinction ratio that can be applied to any wavelength that a gas chamber is not available.

## Result

Figure 2 demonstrates the schematic of the cLSBM system. With the combination of a cylindrical lens C1 and the objective lens, a line beam was generated at the focal plane of the objective lens.

**Figure 2.**
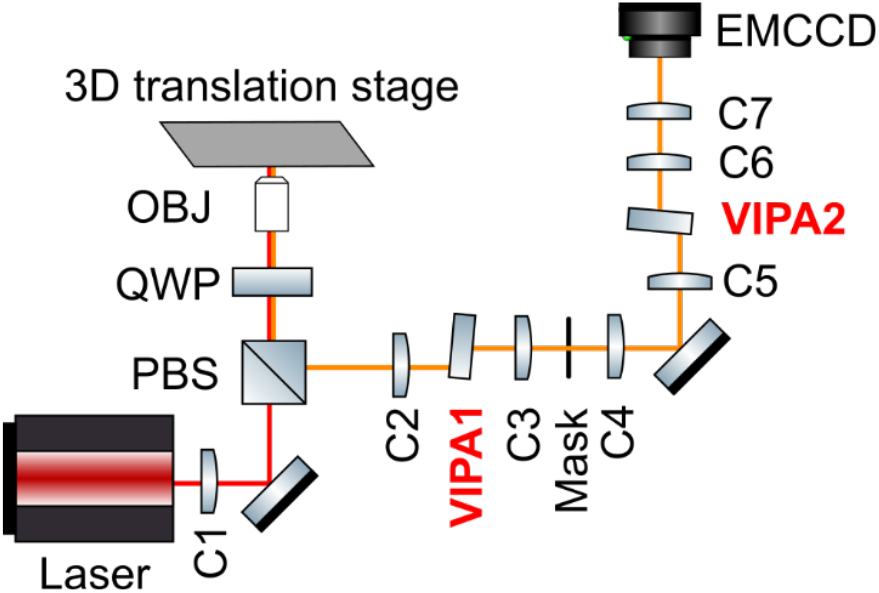
Schematic of the coaxial line-scanning Brillouin microscope. (PBS: polarized beam splitter; QWP: quarter-wave plate; OBJ: objective lens; C1-C7: cylindrical lenses; EMCCD: electron-multiplying charge-coupled device; red line: laser beam path; orange line: Brillouin scattering light path).

The backscattered light was collected by the same objective and projected onto the input window of the first VIPA (VIPA1) through a cylindrical lens (C2). The pattern after VIPA1 was projected onto a mask (Mask) by a cylindrical lens (C3), where the laser noise was rejected by the mask and Brillouin signal passed through Mask (Figure 1(b)). Then, two cylindrical lenses (C4 and C5) were used to realign the Brillouin signal and focused onto the input window of the second VIPA (VIPA2). At last, two cylindrical lenses (C6 and C7) to project the pattern from VIPA2 onto a EMCCD for analysis.

We first used the brightfield camera to capture the illumination beam, and the result showed a line-shaped beam with a size of 153.3×2.7 μm (Figure 3). For the spectrometer part, VIPA1 here was used as a filter to reject the non-Brillouin noise. To quantify the noise rejection of laser components, we first opened the Mask and obtained the energy distribution curve pattern of the laser from the VIPA1 pattern on the Mask (Figure 4(a)). Then, we calculated the area of the laser energy distribution curve under different pixel ranges to mimic the closure of the Mask and the results was shown in Figure 4(b) (blue line). Also, by tuning the window of the Mask, we measured the actual laser suppression at different closure locations (cut-off frequencies), which was shown in Figure 4(b) (red circles). The results showed a laser suppression of ∼-18 dB at the cut-off frequency of 4.9 GHz. Since in 180-degree scattering geometry, the Brillouin shift of the water for 780 nm laser is 5.0 GHz, and most biological sample have higher Brillouin shift than that of water, the Mask won’t reject the Brillouin signal from the sample.

**Figure 3.**
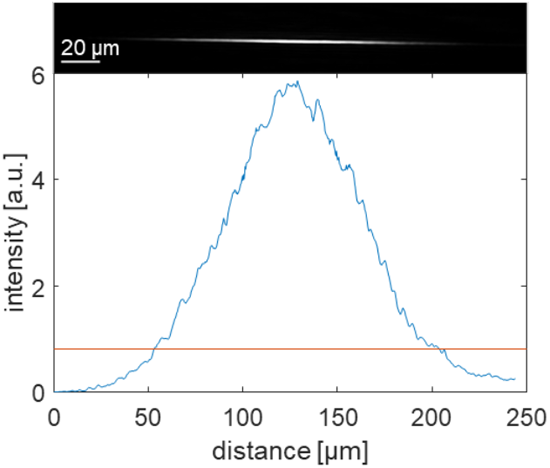
Beam profile of the illumination line at the focal plane of the objective lens.

**Figure 4.**
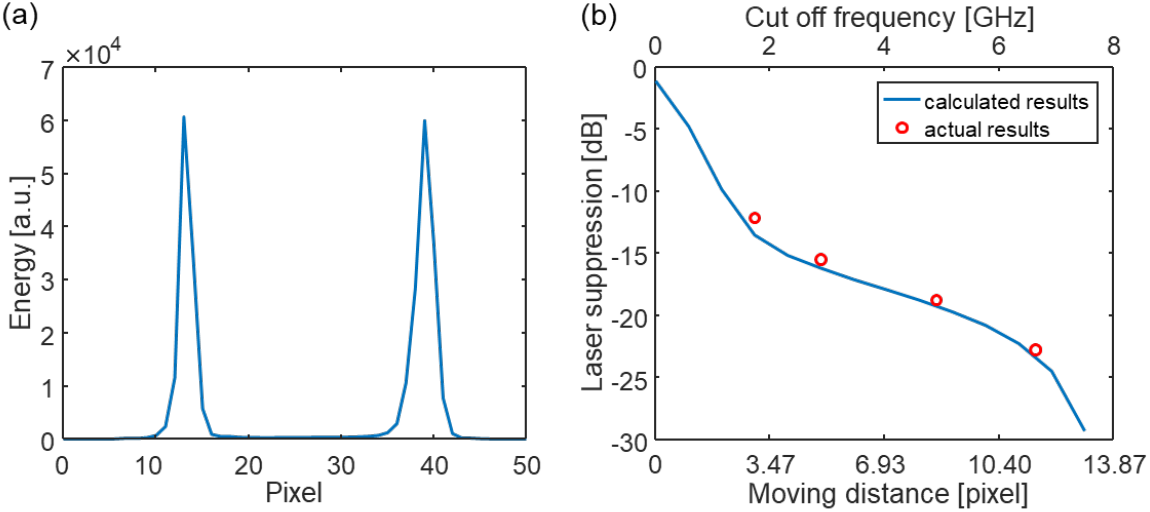
**(a)** the laser distribution curve of the laser from the first VIPA (VIPA1) pattern; **(b)** The laser suppression ability of the Mask when tuning the closure of the mask.

Furthermore, we characterized the performance of the VIPA2 spectrometer. We measured the water and ethanol with the input laser power of 90 mW and camera exposure time of 0.2 s. The Brillouin spectrum of the water and ethanol were shown in Figure 5. We obtained the energy distribution curve at the middle position from the first 6 orders (red dashed line in Figure 5) and calculated the Brillouin shift from Lorentz fitting. The precision was calculated from the standard deviation of 100 repeated measurements and listed in Table 1. Since the Brillouin signal was distributed to multiple orders, to improve precision, we can average the shift results from the 6 orders. Theoretically, this will make a smaller standard deviation following the equation:

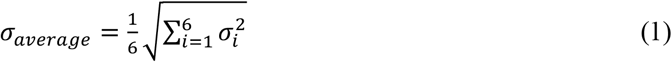

**Figure 5.**
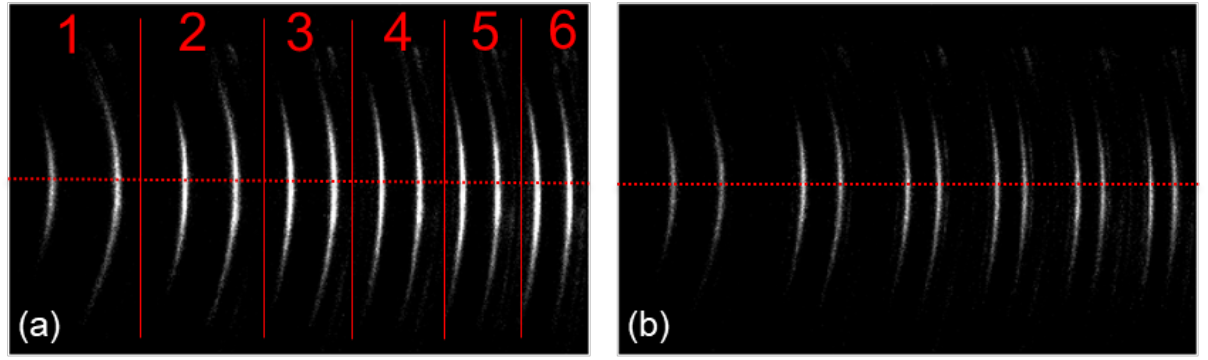
Brillouin spectrum from VIPA2 of **(a)** water, and **(b)** ethanol.

**Table 1.** The precision of the measurement from the first 6 orders.

| Order# | 1 | 2 | 3 | 4 | 5 | 6 |
| --- | --- | --- | --- | --- | --- | --- |
| Ethanol precision [MHz] | 19.2 | 15.3 | 17.0 | 16.5 | 22.5 | 25.7 |
| Water precision [MHz] | 23.2 | 25.4 | 25.6 | 31.2 | 40.8 | 36.3 |

where *σ*_*i*_ is the standard deviation from one single order. The averaged results showed a precision of 7.5 MHz and 12.6 MHz, for ethanol and water respectively, and the results are consistent with theoretical value calculated by the Equation (1) (8.0 MHz and 12.7 MHz, respectively). At last, we measured the SNR of ethanol’s Brillouin signal under different exposure times. We fitted the curve of SNR vs. exposure time on a log-log plot. The slop of the two curves is both close to 0.5, indicating both VIPAs work at shot noise-limited condition (Figure 6).

**Figure 6.**
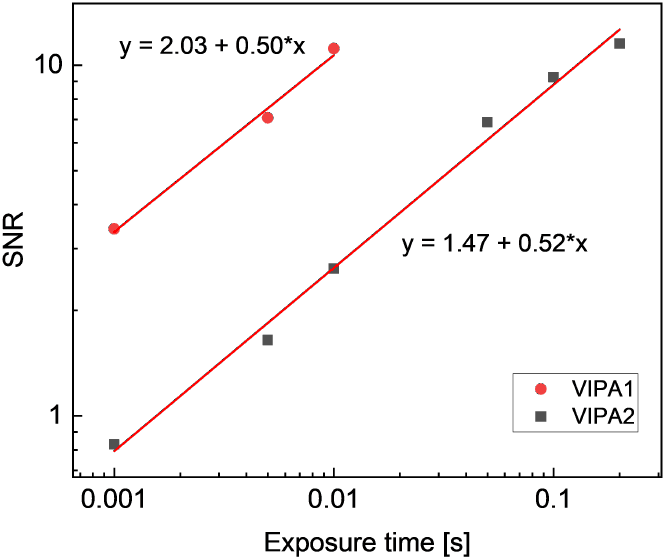
SNR of ethanol signal under different camera exposure times (red circle: SNR from VIPA1; black square: SNR from VIPA2) and the fitting curve of the results (red lines).

To characterize the spatial resolution, we took an image of a polystyrene bead embedded in 1% agarose. By using 1 μm step size along x axis, we scan the central plane of the bead, and the result is shown in Figure 7(b). The diameter of the bead was 50 μm and was confirmed from both bright field image and Brillouin shift image (Figure 7(a) and Figure 7(b)). By moving the bead along y-axis, we characterized the spatial resolution in y-direction (the Brillouin signal line), which shows a spatial resolution of 1.47 μm/pixel and is close to the diffraction limit. We also measure the bead at different depth with a distance of 15 μm, as shown in Figure 7(c). At last, we did a preliminary biological sample measurement using a day-9 tumor spheroid of breast cancer cell MCF10CA1h using the same scanning parameters, as shown in Figure 8. These results demonstrated the availability of the system to be applied to biological tests.

**Figure 7.**
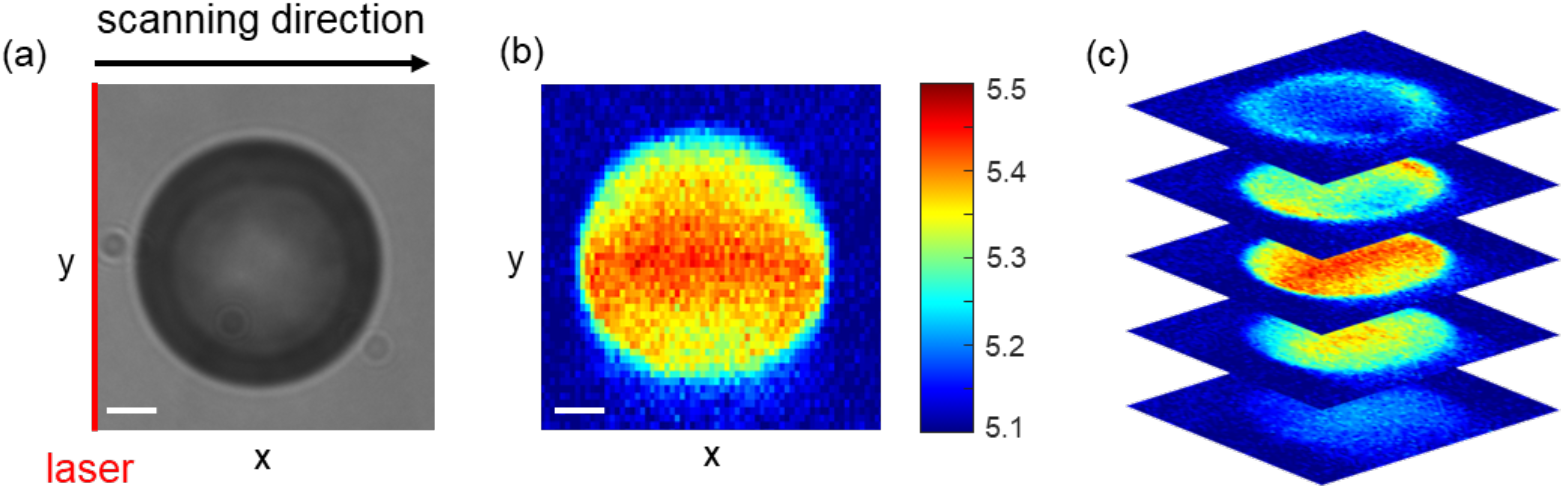
(a) bright-field image of the polystyrene bead; (b) Brillouin shift image of the bead at central plane; (c) Brillouin shift image of the bead at different plane with the step size of 15 μm. (Scale bar: 10 μm)

**Figure 8.**
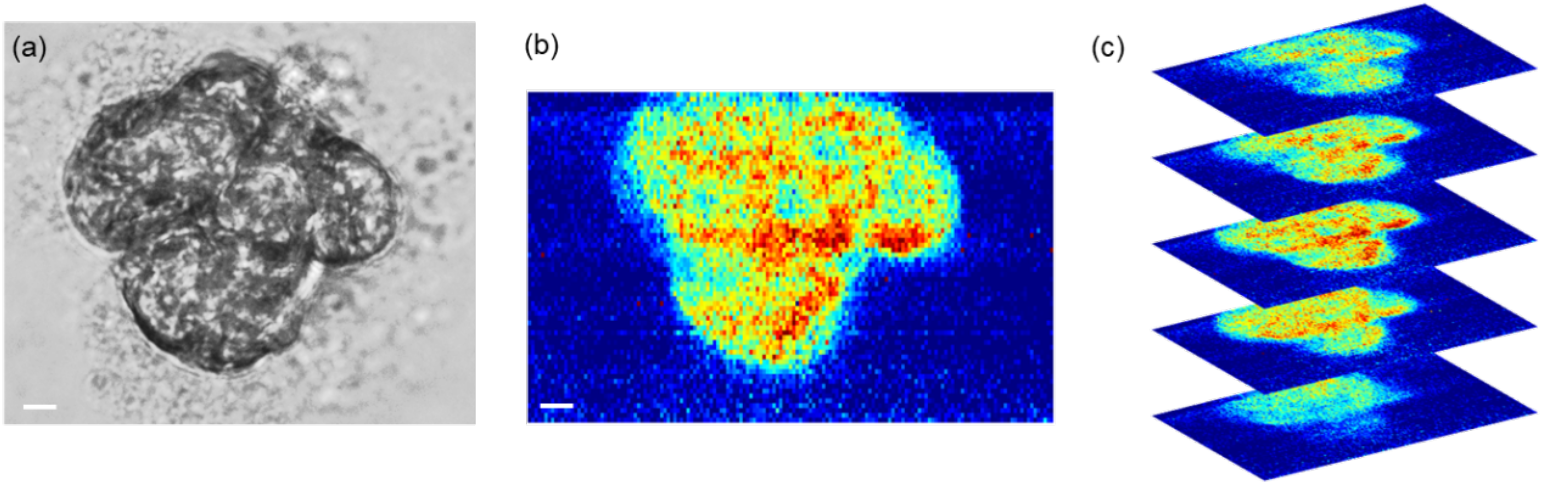
(a) bright-field image of the spheroid; (b) Brillouin shift image of the spheroid at central plane; (c) Brillouin shift image of the spheroid at different plane with the step size of 15 μm. (Scale bar: 10 μm)

## Discussion

In this work, we developed a cLBSM featuring a two-stage parallel VIPA spectrometer design. The inverted microscopic configuration provides easy access to biological samples in standard petri dishes. The two-stage parallel VIPAs achieve a higher extinction ratio, effectively rejecting more non-Brillouin noise and accommodating any laser wavelength.

Currently, the system has two main limitations. First, the laser suppression ability of VIPA1 is relatively low (∼18 dB). Enhancing rejection further would require closing the mask more, but this would also block the Brillouin signal. Second, the signal-to-noise ratio (SNR) for single orders is low. To ensure the two Brillouin peaks are sufficiently close for full coupling into VIPA2, the beam’s convergence angle at the VIPA2 input window must be large, resulting in energy distribution across multiple orders. In the next step, we will optimize the system to address these technical limitations.

